# Ribosomal protein eL22 contributes to the assembly of 60S ribosomal subunits in *Saccharomyces cerevisiae*

**DOI:** 10.64898/2026.05.20.726491

**Authors:** José Fernández-Fernández, Sara Martín-Villanueva, Taylor N. Ayers, Carla V. Galmozzi, John L. Woolford, Jesús de la Cruz

## Abstract

Ribosome biogenesis is a highly coordinated pathway that involves the assembly of ribosomal RNAs (rRNAs) with ribosomal proteins (r-proteins) to generate functional ribosomal subunits (r-subunits). The *Saccharomyces cerevisiae* (yeast) large 60S r-subunit consists of three rRNA molecules and 46 r-proteins. The contributions of nearly all r-proteins of the yeast large r-subunit have been characterized; however, a few non-essential proteins remain poorly understood. Although non-essential, human eL22 has been identified as a key player in p53 regulation during ribosomal stress and as a highly mutated target in cancers. Despite this function, the role of eL22 in ribosome maturation is still ill-defined. In this study, we characterized yeast eL22 r-protein. Our results show that eL22 assembles into intermediate nucleolar pre-60S ribosomal particles. Loss of eL22 impairs cell growth and reduces 60S r-subunit accumulation, phenotypes that are exacerbated at low temperatures. Analysis of pre-rRNA processing by pulse-chase labeling, northern blot hybridization, and primer extension reveals a defect in 27S pre-rRNA maturation, specifically at the level of 27SB pre-rRNA processing. Consequently, nuclear export of eL22-deficient pre-60S particles is mildly impaired. Furthermore, we identify genetic interactions between eL22 and neighboring r-proteins, eL38 and eL31. We conclude that eL22 assembly is required for optimal pre-60S maturation during middle nucleolar stages, particularly at low temperatures, a function likely supported by the cooperative action of other r-proteins associated with common elements of 25S rRNA.

**Highlights:**

- We have studied the role of r-protein eL22 in yeast ribosome assembly.
- eL22 is required for 60S ribosomal subunit production.
- The absence of eL22 is critical at low temperatures.
- eL22 is important for 27SB pre-rRNA processing and nuclear export of pre-ribosomes.
- eL22 functionally interacts with r-proteins eL38 and eL31 in domain III of 25S rRNA.

## Introduction

Ribosomes are ubiquitous ribonucleoprotein complexes that translate mRNAs into proteins [1]. They are present in the cytoplasm of all living cells as well as in the mitochondria and the chloroplasts of eukaryotes. Despite the high degree of evolutionary conservation of their core structure, eukaryotic cytoplasmic ribosomes are the most complex and require the most elaborate biogenesis pathway [2, 3]. For more than 50 years, *Saccharomyces cerevisiae* (hereafter, yeast) has served as a key model organism for the study of ribosome assembly and function (for reviews, see [3-7]). The yeast cytoplasmic ribosome consists of two asymmetric ribosomal subunits (r-subunits). The small 40S r-subunit (SSU), which contains the decoding center, comprises one ribosomal RNA (18S rRNA) and 34 distinct ribosomal proteins (r-proteins). The large 60S r-subunit (LSU), which harbors the peptidyl transferase center and the polypeptide exit tunnel (PET), contains three rRNAs (5S, 5.8S, and 25S rRNA) and 46 r-proteins. High-resolution structures of ribosomes have been determined by X-ray crystallography and cryo-electron microscopy (cryo-EM), revealing the complex architecture of these organelles [2, 6-8].

In yeast, as in all eukaryotes, cytoplasmic ribosome synthesis is a highly coordinated process that proceeds sequentially through the nucleolus, nucleoplasm and cytoplasm [3, 4, 6]. This rapid and highly energy-consuming process supports the production of ca. 2000 ribosomes per minute in exponentially growing cells [9]. In the nucleolus, RNA polymerase I synthesizes a large precursor rRNA (pre-rRNA) that contains the mature 18S, 5.8S and 25S rRNAs, whereas the 5S rRNA is transcribed independently by RNA polymerase III. Pre-rRNAs undergo co-transcriptional covalent modification and pre-rRNA processing (see **Figure S1**). These reactions take place within pre-ribosomal particles, which incorporate the nascent pre-rRNAs, distinct early and late-binding r-proteins, and a large subset of the approximately 300 known assembly factors involved in ribosome maturation [3, 6].

Over the past decades, the structures of numerous assembly intermediates, ranging from the small subunit (SSU) processome to late cytoplasmic pre-SSU and pre-LSU particles, have been structurally characterized at near-atomic resolution by cryo-EM [6]. Analyses of these intermediates have provided key insights into the sequential and irreversible pre-rRNA folding and remodeling events that drive r-subunit maturation, and have clarified the molecular functions of many assembly factors, including their roles in enabling r-protein incorporation [3, 6, 10]. During this dynamic, multistep pathway, early remodeling events promote the release of primary assembly factors, the recruitment of additional factors, and the association of late-binding r-proteins. These coordinated transitions permit continued structural rearrangements within pre-ribosomal particles and propel the maturation of both r-subunits forward [6, 7].

The 25S rRNA consists of six phylogenetically conserved domains (I-VI, 5’ to 3’) of secondary structure that intertwine with each other to form a monolithic tertiary structure [10, 11]. Counterintuitively, assembly of the LSU r-subunits does not strictly proceed in a 5’ to 3’ order; rather, rRNA domains I (including the 5.8S rRNA), II, and VI first assemble during early nucleolar stages with many of their constituent r-proteins to form the solvent-exposed shell of the LSU subunit. At this stage, the middle rRNA domains III, IV, and V remain flexible, then compact onto late nucleolar pre-LSU r-particles to constitute the functional centers on the inter-subunit interface [6, 7]. Domain III rRNA completes the exit platform and a wall of the nascent PET, crucial structures for mediating folding of emerging nascent peptides during protein synthesis. Furthermore, completion of domain III rRNA assembly serves as the catalyst for middle domain maturation and installation of the peptidyl transferase center.

Functional analyses have demonstrated that many r-proteins are active players in the maturation and nucleocytoplasmic transport of pre-ribosomal particles [10-12]. Moreover, cryo-EM studies have provided insights into how r-proteins promote the formation or rearrangement of structures within pre-ribosomal particles. To date, the contributions of virtually all LSU r-proteins to ribosome biogenesis have been examined, with only a few exceptions, including the non-essential eL22, one of nine r-proteins associating to domain III rRNA in the mature LSU (**Figure S2**).

In human cells, eL22 has been identified as a key player in p53 regulation and as a highly mutated target in cancers [13, 14]. Thus, studies have demonstrated that during ribosomal stress, eL22 functions as a tumor suppressor and contributes to activation of p53 and target genes [13]. These findings highlight the critical extra-ribosomal function of eL22; however, its direct role in LSU maturation remains to be defined. In this study, we performed a functional analysis of eL22 during yeast LSU biogenesis. Our data show that the absence of eL22 causes a growth defect that is particularly pronounced at low temperatures. Loss of eL22 also results in reduced LSU levels and a mild impairment in the nuclear export of pre-LSU r-particles. Notably, eL22 is required for efficient processing of 27S pre-rRNAs. Furthermore, our results indicate that eL22 functionally interacts with the adjacent domain III r-proteins eL38 and eL31, suggesting a coordinated role of these r-proteins in the maturation of rRNA domain III or downstream assembly stages.

## Results

### eL22 assembles with pre-LSU particles during middle-nucleolar stages of assembly

The r-protein eL22 is a small (ca. 13.7 kDa), predominantly globular, eukaryote-specific LSU protein located near the exit of the PET. It is embedded between eL31, and the globular domains of eL19 and eL38, and establishes contacts with the external surface of domain III of the 25S rRNA (**Figure S2**). Assembly of most r-proteins occurs in the nucle(ol)us, although a few LSU proteins appear to stably load only with cytoplasmic pre-LSU r-particles [10, 11]. In the nucle(ol)us, r-proteins can bind pre-ribosomal intermediates at different stages of assembly, ranging from early to late intermediates [10, 11, 15, 16]. The precise timing of eL22 assembly remains unclear; cryo-EM analyses indicate that eL22 is among the r-proteins present in late nucleolar pre-LSU r-particles (state D), in which domain III of 25S rRNA is clearly visible [17]. However, whether eL22 associates with pre-LSU assembly intermediates at an earlier stage of biogenesis when domain III is not yet visible by cryo-EM has not yet been resolved.

To investigate the timing of eL22 assembly, we expressed an eL22A protein tagged at its C-terminus with yeast-enhanced GFP (yEGFP) from its native promoter on a *CEN* plasmid. The resulting eL22-yEGFP fusion protein was fully functional, as its expression complemented both the growth and ribosome biogenesis defects (see below) of an *rpl22* null strain at 30 °C (**Figure S3**). We next monitored the localization of eL22-yEGFP upon expression of a dominant-negative Nmd3-Δ100 variant under the control of an inducible *GAL* promoter. This truncated variant lacks the nuclear export signal (NES) of Nmd3 and consequently prevents pre-LSU export, causing the accumulation of particles in the nucle(ol)us [18]. Thus, if eL22 assembles in the nucle(ol)us, we would expect nuclear accumulation of eL22-yEGFP signal. Under *NMD3-Δ100-*inducing conditions, eL22-yEGFP accumulated in the nucleus in more than 80% of the examined cells. In contrast, under non-inducing conditions, consistent with its role as a r-protein, eL22-yEGFP was predominantly detected in the cytoplasm (**Figure S4**). As an alternative approach to investigate when eL22 assembles with pre-ribosomes, we assayed with which consecutive pre-rRNA processing intermediates eL22 copurified. To do so, we affinity-purified particles containing eL22-yEGFP using GFP-Trap beads and analyzed the associated pre-rRNAs by northern blot hybridization (see **Figure S1** for a schematic of the pre-rRNA processing pathway). As shown in **Figure 1**, eL22-yEGFP stably co-purified with 27SB pre-rRNAs and downstream intermediates, such as the 7S pre-rRNAs. As expected for a r-protein, substantial amounts of mature rRNAs also co-purified. In contrast, we detected no significant amounts of the 27SA_2_ pre-rRNA, which is the major intermediate present in the earliest pre-LSU particles, were detected. Consistent with these findings, eL22 was substantially enriched in association with nucleolar and nucleoplasmic pre-LSU particles affinity purified using TAP-tagged Nop7 and Arx1 (T.A. and J. L.W., unpublished results). Taken together, these results strongly suggest that eL22 stably associates with middle nucleolar pre-LSU particles, but not with earlier intermediates.

**Figure 1.**
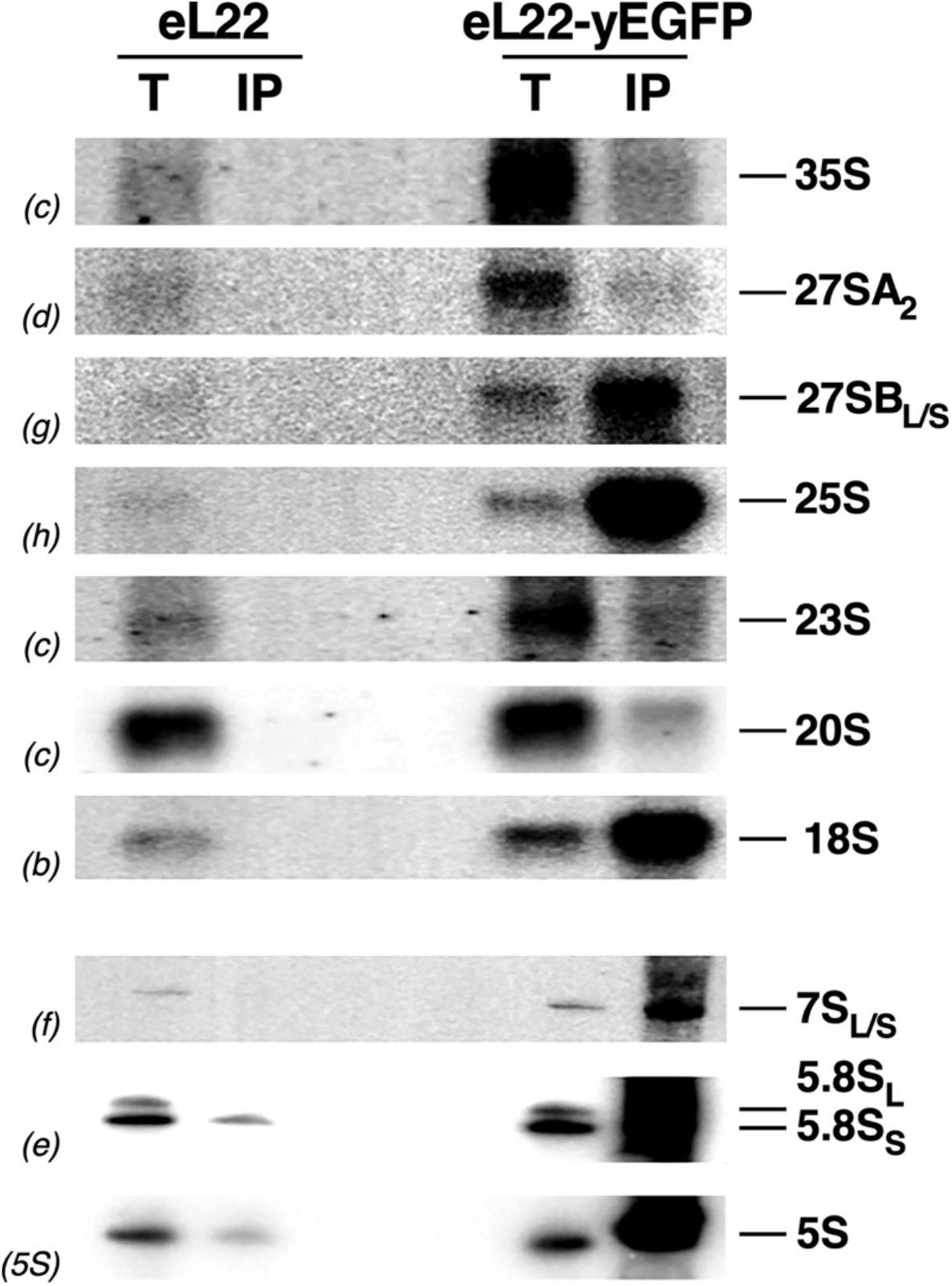
eL22-yEGFP stably associates with pre-LSU particles containing 27SB pre-rRNAs. Cells of the *rpl22* null strain expressing either untagged eL22 or yEGFP-tagged eL22 from a plasmid were grown in liquid SD-Leu medium at 30 °C to mid-log phase. Whole-cell extracts were prepared, and r-particles were affinity-purified using the GFP-Trap immunoprecipitation method. Total RNA was extracted from whole cell extracts (T) and immunoprecipitates (IP) and analyzed by northern blotting. Probes used to detect the different pre- and mature rRNAs are shown in parentheses and are described in **Table S3**.

### Loss of eL22 results in a mild growth defect that is exacerbated at low temperatures

Like most yeast r-proteins [19-21], eL22 is encoded by two paralogous genes, *RPL22A* (*YLR061W*) and *RPL22B* (*YEL034C-A*), which share 84.6% identity and 94.3% similarity [19]. To investigate the role of eL22 in yeast ribosome biogenesis, we first analyzed the growth phenotypes of strains lacking either *RPL22A* (hereafter, *rpl22AΔ* strain), *RPL22B* (hereafter, *rpl22BΔ* strain), or both genes simultaneously (hereafter, *rpl22Δ* or *rpl22* null strain). As shown in **Figure 2**, cells lacking eL22 displayed a noticeable slow-growth phenotype at 30 °C. This defect was almost completely suppressed at 37 °C but was markedly exacerbated at lower temperatures (22 °C). Importantly, deletion of *RPL22A* was responsible for this growth defect, as the *rpl22* null strain exhibited growth defects similar to those of the *rpl22AΔ* mutant, whereas the *rpl22BΔ* mutant grew indistinguishably from the isogenic wild-type strain at all temperatures tested (**Figure 2)**. The lack of a growth defect in the *rpl22BΔ* mutant likely results from *RPL22B* being expressed at much lower levels than *RPL22A*, as is sometimes the case for yeast r-protein paralogous genes [22]. These results are consistent with previously published observations and indicate that eL22 is required for optimal cellular growth, particularly under low-temperature conditions [23, 24].

**Figure 2.**
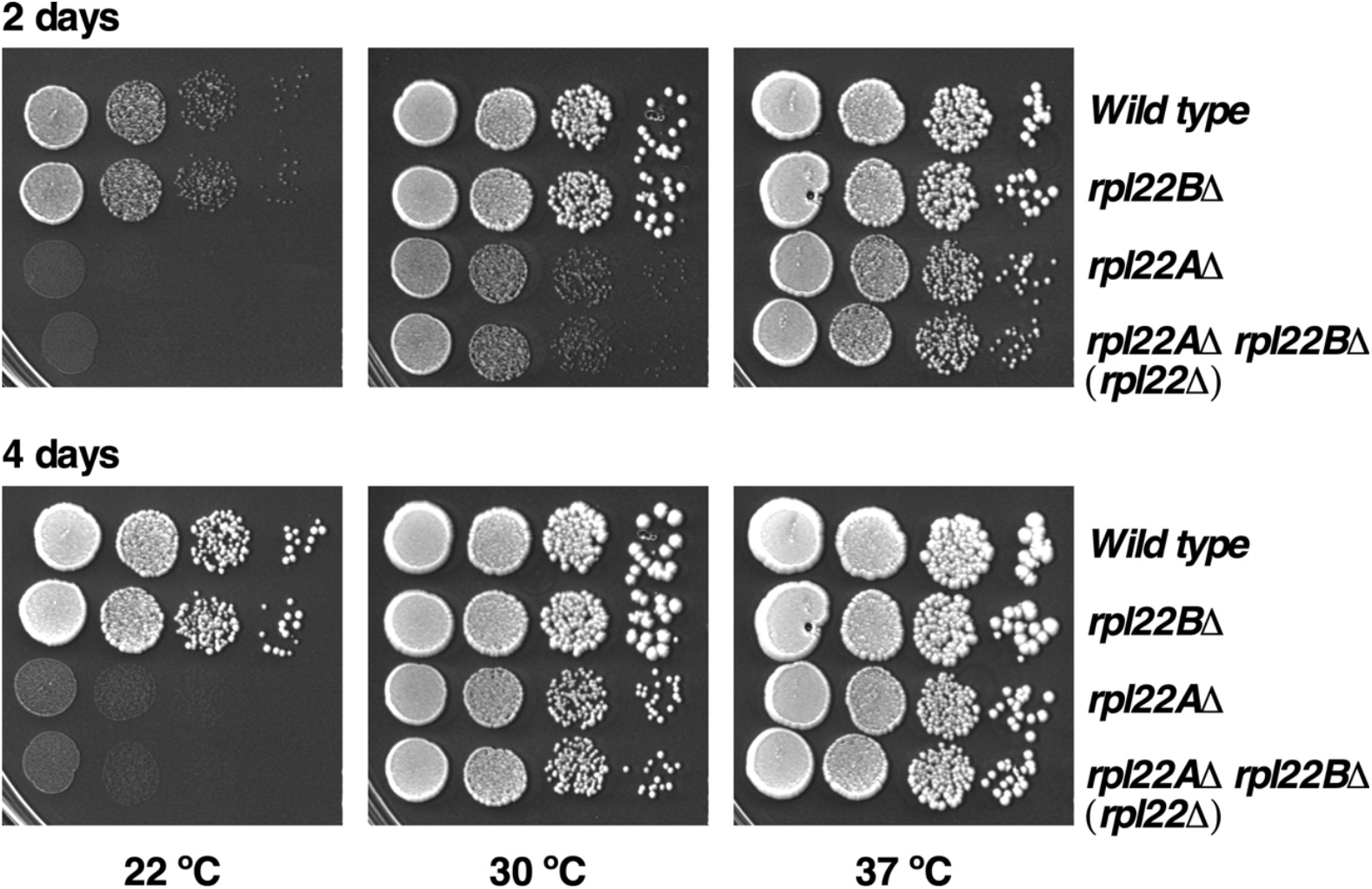
Cells lacking eL22 exhibit a cold-sensitive growth phenotype. Growth comparison of isogenic wild-type, single *rpl22AΔ* and *rpl22BΔ*, and *rpl22Δ* null strains. Cells were grown in liquid YPAD medium at 30 °C to mid-log phase, diluted to an OD_600_ of 0.05, and 10-fold serial dilutions were spotted onto YPAD plates. Plates were incubated at 22 °C, 30 °C and 37 °C for 2 days (upper panels) and 4 days (lower panels).

### eL22 is required for normal LSU accumulation

To determine whether the absence of eL22 affects the production of mature LSUs, we analyzed polysome profiles from the *rpl22AΔ* and *rpl22BΔ* single mutants, as well as from the *rpl22* null strain, grown at 30 °C. As shown in **Figure 3**, both *rpl22AΔ* and *rpl22Δ* mutants exhibited similar aberrant polysome profiles compared with the isogenic wild-type strain. These defects were characterized by a clear decrease in the levels of free LSUs relative to free SSUs and by the presence of half-mer polysomes. Such profiles are a hallmark of mutants specifically impaired in LSU biogenesis (e.g., [25-27]). In contrast, consistent with its wild-type-like growth phenotype, *rpl22BΔ* cells displayed a normal polysome profile, similar from that of the isogenic wild-type strain. Quantification of total r-subunits by low-Mg^2+^ sucrose gradients further revealed that loss of eL22 results in an approximately 35% specific reduction in LSUs relative to SSUs compared with the wild-type control (data not shown).

**Figure 3.**
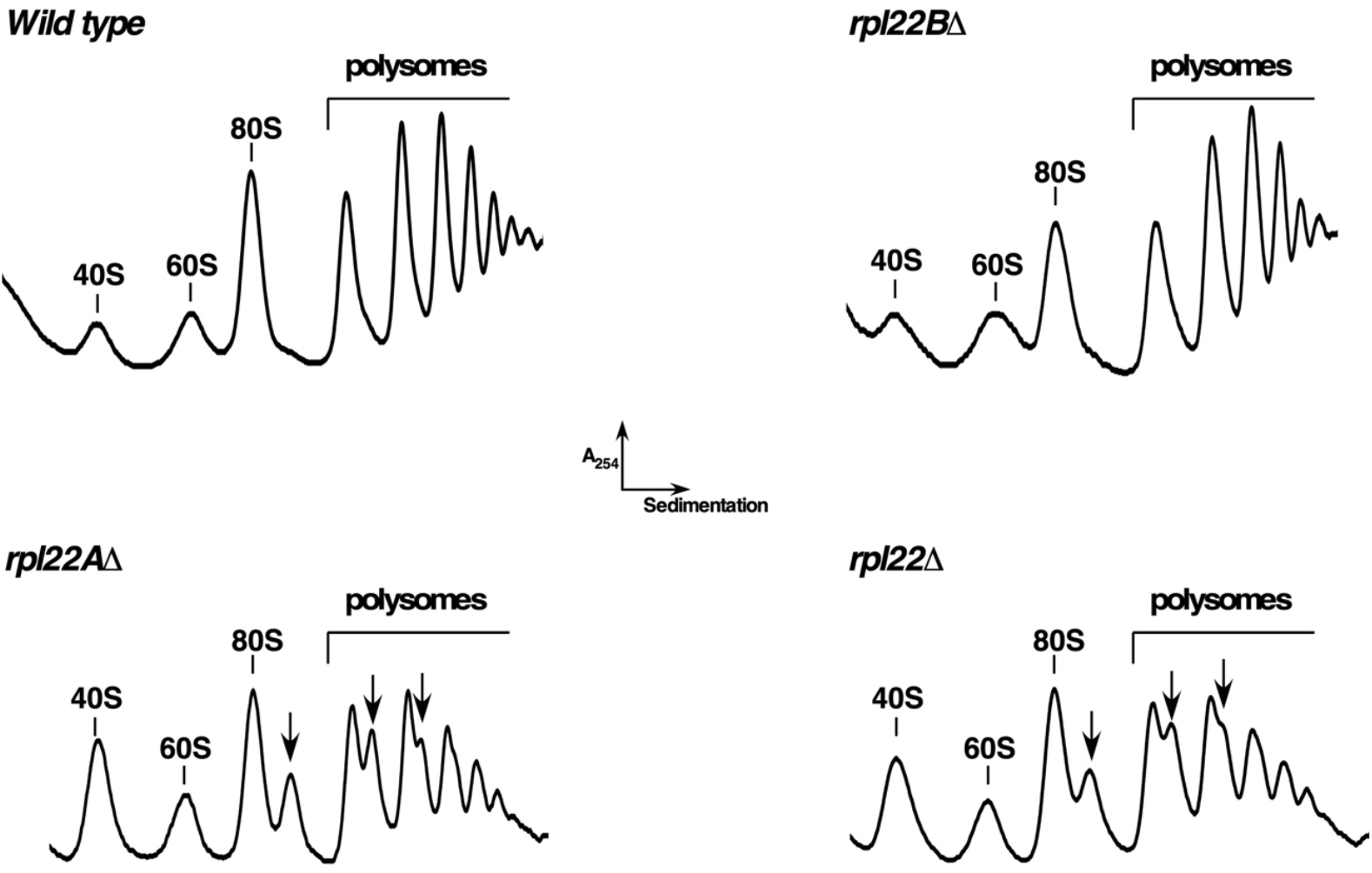
Loss of eL22 results in a deficit of LSUs. Polysome profile analysis of isogenic wild-type, single *rpl22AΔ* and *rpl22BΔ*, and *rpl22Δ* null strains. Cells were grown in liquid YPAD medium at 30 °C to mid-log phase. Whole-cell extracts were prepared, and 10 A_260_ units of each extract were resolved in 7-50% sucrose gradients. The A_254_ was continuously recorded. Sedimentation is from left to right. Peaks corresponding to free SSUs (40S) and LSUs (60S), vacant ribosomes/monosomes (80S), and polysomes are indicated. Half-mer polysomes are marked with arrows.

We also examined how the absence of eL22 affects LSU production at 22 and 37 °C. Interestingly, the severity of the LSU defect slightly increases when *rpl22* null cells were grown at 22 °C (**Figure S5A**), but it was almost completely suppressed when cells lacking eL22 were cultured at 37 °C (**Figure S5B**). We therefore conclude that eL22 plays an important role in LSU production, particularly at low and normal temperatures. We speculate that eL22 functions in ribosome assembly by promoting proper pre-rRNA folding or through its interaction with a critical assembly factor, steps that may be especially susceptible to cold-induced perturbations that ultimately impair cell growth (for discussion, see [28]).

### eL22 is required for normal 27SB pre-rRNA processing

To determine whether the reduction in LSU levels observed in the absence of eL22 results from an impairment of a specific step of the pre-rRNA processing pathway, we analyzed the steady-state levels of pre- and mature rRNAs in single and double *rpl22* mutants by northern blot hybridization and primer extension analyses. To do this, we first extracted and analyzed RNA from wild-type cells, *rpl22AΔ* and *rpl22BΔ* single mutants, and the *rpl22* null strain grown in YPAD at 30 °C. Consistent with the polysome profile data, no detectable defects were observed in the *rpl22BΔ* mutant relative to the isogenic wild-type strain (**Figure 4**). In contrast, the *rpl22AΔ* mutant and the *rpl22* null strain displayed mild but comparable defects in pre-rRNA processing. These mutants showed a slight accumulation of both 35S and 23S pre-rRNAs (**Figure 4A**), a phenotype commonly reported in many other mutants related to LSU biogenesis (see Discussion). We also detected a significant accumulation of 27SB pre-rRNA together with a modest decrease in mature 25S rRNA (**Figure 4A**). However, the increase in 27SB pre-rRNAs was not accompanied by detectable changes in the steady-state levels of 7S pre-rRNAs or mature 5.8S and 5S rRNAs (**Figure 4B**). Primer extension confirmed elevated levels of both 27SB_L_ and 27SB_S_ pre-rRNAs, whereas 27SA_2_ levels remained essentially unchanged and 27SA_3_ levels showed a slight increase (**Figure 4C**). To assess whether the growth and polysome defects of the *rpl22* null strain correlate with an impairment in pre-rRNA processing, we also performed northern blot analyses on cells grown at 22 and 37 °C. Strikingly, as shown in **Figure S6**, while the accumulation of 27SB pre-rRNAs persisted at 22 °C, nearly wild-type levels of these and other precursors (e.g. 35S pre-rRNA) were detected at 37 °C.

**Figure 4.**
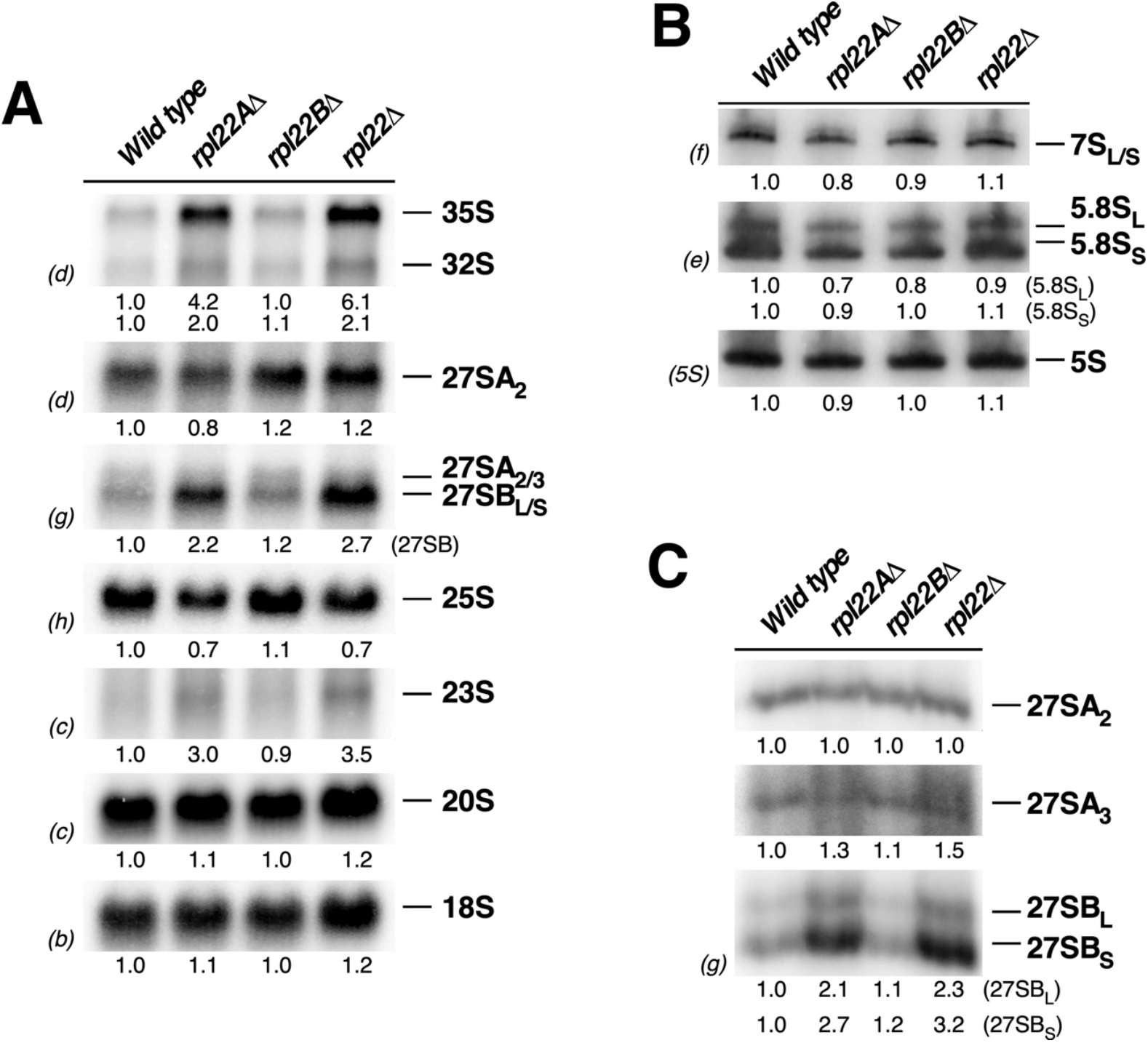
Absence of eL22 impairs 27SB pre-rRNA processing. Isogenic wild-type, single *rpl22AΔ* and *rpl22BΔ*, and *rpl22Δ* null strains were grown in liquid YPAD medium at 30 °C to mid-log phase. Total RNA was extracted, and equal amounts (5 µg) were analyzed by northern blotting or primer extension. (**A**) Northern blot analysis of high-molecular-mass pre- and mature rRNAs. (**B**) Northern blot analysis of low-molecular-mass pre- and mature rRNAs. (**C**) Primer extension analysis using probe g as the primer. Signal intensities were quantified by phosphorimager scanning (values shown below each panel) and normalized to wild-type levels, which were arbitrarily set to 1.0. Probes used are indicated in parentheses and described in **Table S3**.

To further ascertain the contribution of eL22 to the dynamic conversion of pre-rRNAs into mature rRNAs, we examined the kinetics of rRNA formation by [5,6-^3^H]uracil pulse-chase analysis. For this purpose, the *rpl22* null strain and its isogenic wild-type counterpart were grown at 30 °C in SD-Ura. As a result, both strains exhibited very similar overall kinetics of rRNA production; however, *rpl22* null cells displayed a mild delay in the conversion of 27SB pre-rRNAs into mature 25S rRNA, as well as a slight delay in the conversion of 7S pre-rRNAs into mature 5.8S rRNAs (**Figure 5**). However, these delays had only minor consequences for the efficiency of accumulation of the corresponding mature 25S and 5.8S rRNAs. No apparent impairment was observed in the processing of 20S pre-rRNA into mature 18S rRNA or the synthesis of 5S rRNA.

**Figure 5.**
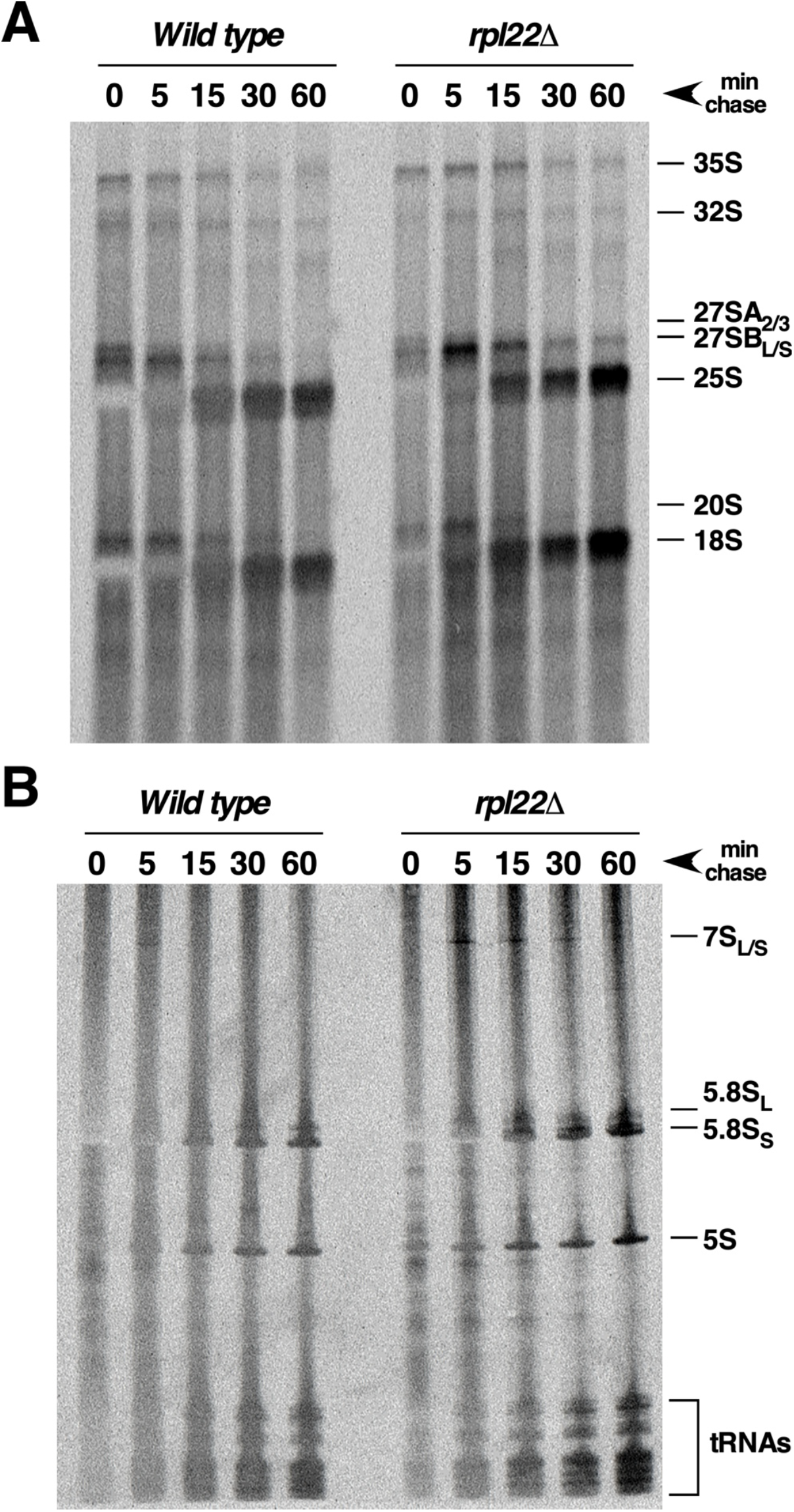
Loss of eL22 delays processing of 27S and 7S pre-rRNAs. Wild-type and *rpl22Δ* null strains were transformed with an empty YCplac33 (*CEN URA3*) plasmid and grown in liquid SD-Ura medium at 30 °C to mid-log phase. Cells were pulse-labeled for 2 min with [5,6-^3^H] uracil and chased for 5 to 60 min with excess unlabeled uracil. Total RNA was extracted, and approximately 3,000 cpm per sample were separated on (**A**) 1.2% agarose-6% formaldehyde gels or (**B**) 7% polyacrylamide-8 M urea gels. RNA was transferred to nylon membranes, exposed to a ^3^H-sensitive phosphorimager screen, and analyzed by phosphorimager scanning. Positions of the indicated pre- and mature rRNAs are shown.

Altogether, these results indicate that eL22 plays a modest but measurable role in rRNA maturation. Its absence causes only a minor defect in the specific branch of the pre-rRNA processing pathway that decreases the efficiency of conversion of 27SB pre-rRNAs into mature 25S and 5.8S rRNAs. Notably, this defect is significantly suppressed when *rpl22* null cells are grown at elevated temperatures.

### The absence of eL22 leads to a mild defect in the nuclear export of pre-LSU particles

To further assess the consequences of the absence of eL22 on LSU maturation, we examined whether the *rpl22Δ* mutant was impaired in the nuclear export of pre-LSU particles. For this purpose, we analyzed the localization of uL23-yEGFP, which was used as an LSU reporter, in *rpl22* null and isogenic wild-type cells grown at 22 °C. As shown in **Figure S7**, the steady-state levels of uL23-yEGFP were cytoplasmic in wild-type cells, as expected for an r-protein. In contrast, a mild nuclear retention of the fluorescent signal of the uL23-yEGFP reporter was observed in some cells (ca. 30%) of the *rpl22* null strain. Comparison with the localization of the nucleolar reporter mRFP-Nop1 indicated that this retention occurred mainly within the nucleolus. This phenotype was specific to the LSU reporter, as when we analyzed the localization of uS3-yeGFP, which was used as an SSU reporter, it localized to the cytoplasm in both wild-type and *rpl22* null strains. Only very subtle differences in the localization of uL23-yEGFP were observed between wild-type and *rpl22* null cells grown at 30 °C (data not shown). Because a direct role for eL22 nucleo-cytoplasmic export of pre-LSU particles is unlikely, we infer that this defect arises from a partial block in late nucleolar or nucleoplasmic stages of LSU maturation, preventing recruitment of nuclear export factors [29]. Similar conclusions have previously been reported upon the deletion, depletion or mutation of numerous other LSU protein genes (see Discussion).

### Functional interactions of eL22 with its neighboring eL31, eL19 and eL38 r-proteins

As noted above, eL22 is an r-protein that contacts rRNA domain III in mature LSUs, where it is positioned adjacent to other r-proteins, including eL31 and eL38 (**Figure S2**). Notably, as eL22, eL31, and eL38 are also mostly globular, and non-essential r-proteins [8, 23] that assemble during intermediate steps of LSU maturation [10, 17], likely in a coordinated manner as rRNA domain III undergoes compaction (discussed in [10]). To begin exploring this issue, we first tested whether the growth defect of the *rpl22* null strain is exacerbated by reducing the cellular pool of eL31 or eL38, as well as that of eL19, another r-protein associated with rRNA domain III (**Figure S2**).

In yeast, loss of eL38, which is encoded by a single gene, does not cause detectable growth impairment ([23] and **Figure 6**). In contrast, the absence of eL31, which is encoded by two paralogous genes, results in a severe growth defect largely attributable to the *RPL31A* paralogue ([23] and **Figure 6**). eL19 is an essential r-protein encoded by two paralogous genes, and in this case, the *rpl19BΔ* mutant displays a pronounced slow-growth phenotype ([23] and **Figure 6**). As shown in **Figure 6**, the introduction of either *rpl31AΔ* or *rpl38Δ* alleles into the *rpl22* null background markedly enhanced the growth defect of the *rpl22* mutant, particularly at 22 ºC. This result suggests that eL31 and eL38 cooperate with eL22 to facilitate optimal maturation, likely by mediating proper folding of domain III rRNA. In contrast, the *rpl19BΔ* allele did not exacerbate the growth impairment of the *rpl22* null strain. Unlike eL31 and eL38, eL19 is located on the opposing interface from eL22, indicating that eL19 may serve in an alternate function from these proteins. Moreover, the growth defect of the double *rpl22Δ rpl38Δ* mutant was more evident when cells were treated with sublethal concentration of specific translation antibiotics (**Figure S8**). These results support the existence of a specific functional interaction between eL22 and its neighboring r-proteins eL31 and eL38, likely reflecting their coordination roles during rRNA folding and LSU assembly and/or translation.

**Figure 6.**
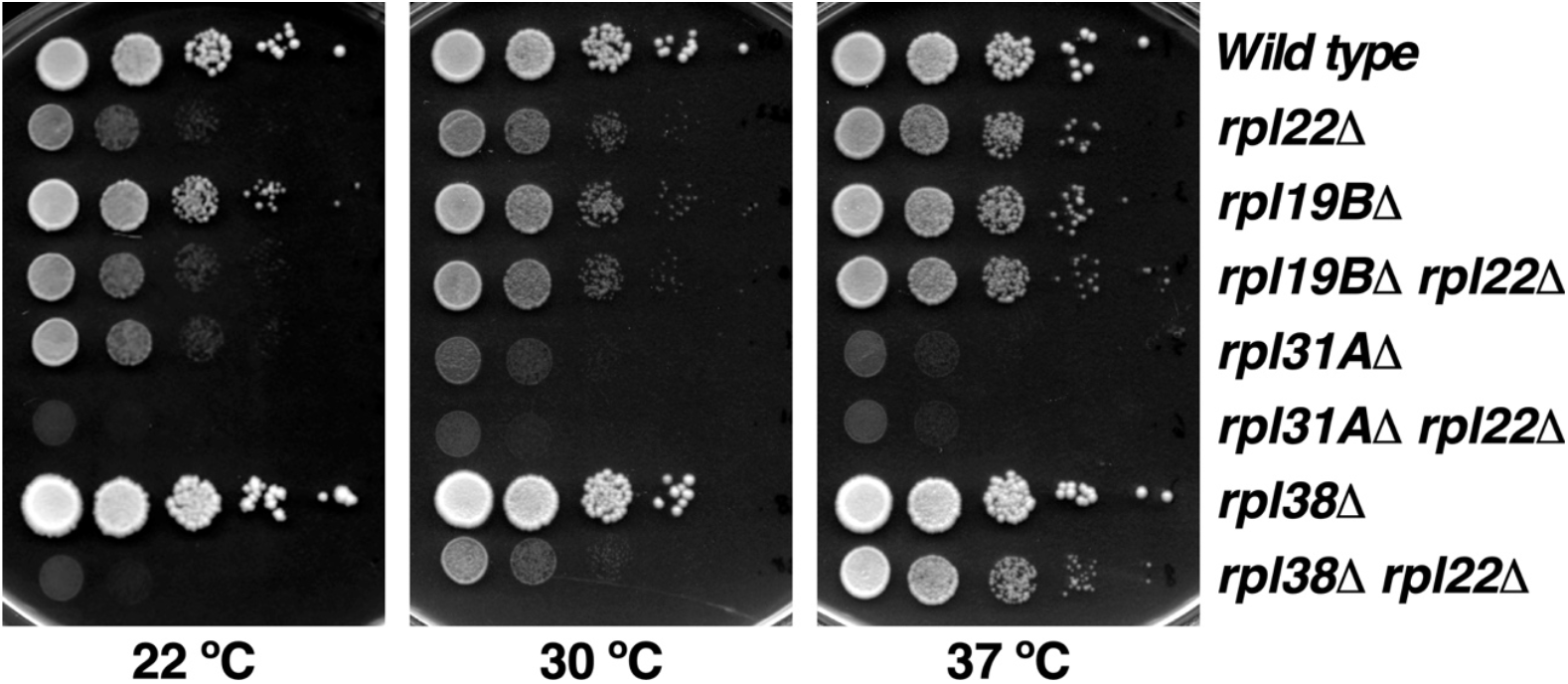
eL22 functionally interacts with eL31 and eL38. Growth assays of strains lacking eL22 in combination with deletions of *RPL19B, RPL31A* or *RPL38*. Wild-type, single-, double-, and triple-deletion strains were grown in YPAD medium at 30 °C to mid-log phase, diluted to an OD_600_ of 0.05, and 10-fold serial dilutions were spotted onto YPAD plates. Plates were incubated at 22 °C for 4 days and at 30 ºC and 37 °C for 2 days. Note that only deletion of *RPL31A* or *RPL38* enhances the slow-growth phenotype of *rpl22* null cells, particularly at 22 °C.

## Discussion

In this work, we have addressed the role of the yeast r-protein eL22 in ribosome biogenesis. eL22 is an evolutionarily conserved protein in eukaryotes that has been identified in only a few archaea belonging to the *Asgard* phylum [30].

In yeast, eL22 is a non-essential r-protein [23]. Cryo-EM studies have detected eL22 in middle nucleolar pre-LSU r-particles (state D) [17], a stage in which 25S rRNA domain III is already compacted. Consistent with these observations, our biochemical analyses show that eL22 stably co-purifies with 27SB-containing pre-LSU particles, whereas it associates only weakly with earlier 27SA-containing intermediates. Yet, these data do not clarify whether eL22 is initially recruited at earlier nucleolar stages to facilitate the construction and compaction of rRNA domain III. Depletion of essential r-proteins that result in an early block of LSU maturation did not impair the association of eL22 with purified pre-LSU particles [16, 31-33], suggesting that eL22 may assemble with domain III at earlier nucleolar stages. Indeed, it has been proposed that most LSU r-proteins bind with pre-LSU r-particles at very early stages with low affinity and only become stably assembled at later steps of maturation [15].

Herewith, we have analyzed the consequences of eL22 absence on ribosome biogenesis. Loss of eL22 causes a mild deficit in LSU production, which likely underlies the observed growth defect of eL22-deficient cells. Processing of the 27SB_L/S_ pre-rRNAs is delayed, and production of mature 25S is slightly reduced. As commonly observed in mutants with impaired LSU biogenesis, 35S and 23S pre-rRNAs accumulate as a result of delayed A_0_-A_2_ site processing. These defects are likely to generate improperly or incompletely assembled pre-LSU r-particles whose nucleo-cytoplasmic export is impaired, leading to mild nucle(ol)ar retention of eL22-deficient pre-LSU r-particles, which we estimate to represent approximately 30% of the total population. Similar phenotypes have been reported for numerous other LSU biogenesis mutants, including mutants of LSU r-protein genes [11]. These effects are generally interpreted as a block in r-subunit maturation before the formation of nuclear export factor binding sites for nuclear export factors and/or the activation of a nuclear quality-control mechanism that eliminates structurally incomplete intermediates, rather than the direct defect in the export machinery [29, 34].

Notably, accumulation of 27SB pre-rRNA, delayed 27SB pre-rRNA processing and nuclear retention of pre-LSU r-particles have also been reported upon depletion or loss-of-function mutation of other domain III r-proteins, including eL19, uL23, eL31 and eL34 [35-37]. This observation suggests that proper formation of 25S rRNA domain III is required for efficient ITS2 cleavage. Alternatively, or in addition, the absence of eL22 might impair the recruitment of *trans*-acting factors required for 27SB pre-rRNA processing. In support of this idea, assembly of essential r-proteins of domain III such as uL23 and eL19 is required for efficient recruitment of assembly factors such as Rsa4, Nog2 and Nop53, all of which are necessary for 27SB pre-rRNA processing [15, 38]. One possible candidate whose recruitment could be affected in the absence of eL22 is Nog1, as it is required for 27SB pre-rRNA processing [39] and displays close physical interactions with eL22 and other r-proteins from rRNA domain III in late pre-LSU r-particles [40]. However, neither the association of Nog1 with pre-LSU r-particles nor the growth defect of several *nog1* alleles (kindly provided by V. G. Panse [41]) are compromised in the absence of eL22 (our unpublished results).

A key finding of this study is that the growth and ribosome biogenesis defects of *rpl22Δ* cells are strongly temperature dependent: they are exacerbated at 22 ºC and largely suppressed at 37 ºC. The accumulation of 27SB pre-rRNA exclusively at low temperatures suggests the presence of a kinetic bottleneck in ITS2 processing. One plausible explanation is that eL22 functions as a structural scaffold that stabilizes a specific pre-rRNA conformation required for efficient processing; in its absence, the rRNA may misfold into off-pathway intermediates that are not resolved, and become rate limiting only at low temperatures, but can be bypassed at higher temperatures. Panse and co-workers have recently demonstrated that the assembly factor Puf6 lowers the activation energy necessary for formation of a specific tertiary structure between distinct 25S rRNA helices [28]. Like *rpl22Δ, puf6* mutants exhibit growth and ribosome biogenesis defects predominantly at low temperatures [28]. Interestingly, yeast cells globally upregulate ribosome assembly factors and r-proteins at low temperatures, and eL22 itself is among the r-proteins most strongly induced at under these conditions [28].

Finally, we examined the functional relationship between eL22 and its neighboring r-proteins eL31 and eL38 in the mature LSU. The more severe growth defect observed in *rpl22Δ rpl38Δ* and *rpl22Δ rpl31AΔ* double mutants compared with the corresponding single mutants strongly suggest that these proteins act cooperatively during ribosome biogenesis and/or function. Consistent with this model, the *rpl22Δ rpl38Δ* double mutant shows increased sensitivity to translation inhibitors such as neomycin and cycloheximide, which also perturb ribosome biogenesis, (e.g. [35, 42-44]). In this context, eL22 and eL38 may cooperate to optimize the architecture of a specific rRNA module within domain III during nucleolar stages of assembly. Although eL19 is also located in proximity to eL22, loss of eL19B does not display comparable synthetic interactions with eL22, suggesting that eL19 does not participate in a shared function.

In conclusion, unlike many essential r-proteins, eL22 represents a eukaryote-specific evolutionary addition to the LSU. eL22 appears to enhance the efficiency and robustness of ribosome biogenesis, particularly under suboptimal growth conditions. We suspect that eL22 promotes formation of rRNA structures in domain III rRNA. Further studies will be required to elucidate the precise molecular mechanisms underlying these LSU assembly defects in the absence of eL22.

## Material and Methods

### Strains and microbiological methods

All *S. cerevisiae* strains used in this study are listed in **Table S1**. Strains JFY010 (*rpl22A*::kanMX4), JFY011 (*rpl22B*::kanMX4), JFY084 (*rpl38*::kanMX4), JFY449 (*rpl31A*::kanMX4), and SMY714 (*rpl19B*::kanMX4) are meiotic segregants derived from the corresponding heterozygous diploid strains obtained from the Euroscarf collection. To generate double and triple deletion mutants, the appropriate strains were crossed, the generated diploids were induced to sporulate, and tetrads were dissected. Strain JFY012 *(rpl22A*::kanMX4 *rpl22B*::kanMX4) is a haploid segregant isolated from the cross between JFY010 and JFY011. Strains JFY114 (*rpl22A*::kanMX4 *rpl22B*::kanMX4 *rpl38*::kanMX4), JFY531 (*rpl22A*::kanMX4 *rpl22B*::kanMX4 *rpl31A*::kanMX4), and SMY712 (*rpl22A*::kanMX4 *rpl22B*::kanMX4 *rpl19B*::kanMX4) were obtained as haploid segregants after crossing JFY012 with JFY084, JFY449, and SMY714, respectively. The correctness of all deletions was confirmed by PCR.

Yeast strains were routinely cultured at the permissive temperature of 30 °C. When indicated, cells were grown at 22 °C or 37 °C. Cultures were maintained either in rich YPAD medium (1% yeast extract, 2% glucose, 2% peptone, 0.2% adenine) or in synthetic minimal SD medium (0.5% ammonium sulphate, 0.15% yeast nitrogen base, 2% glucose) supplemented with the appropriate amino acids and bases. When required, 2% glucose was replaced with 2% galactose as the carbon source (SGal medium). Solid media were prepared by adding 2% agar. Growth and handling of yeast, and media preparation were performed according to standard procedures [45]. When necessary, cells were transformed with the appropriate plasmids using the lithium acetate method [46]. Tetrads were dissected using a Singer MSM200 micromanipulator.

### Plasmids and oligonucleotides

The *Escherichia coli* DH5α strain was used for cloning and propagation of all plasmid vectors employed in this study. A complete list of plasmids is provided in **Table S2**. DNA manipulation procedures were carried out according to standard protocols [47]. Construction of the plasmids YCplac111-RPL22A-HA and YCplac111-RPL22A-GFP was achieved by PCR amplification of the corresponding DNA fragments using specific oligonucleotides. The resulting PCR products were subsequently cloned into the appropriate vectors and verified by DNA sequencing. Additional details regarding cloning strategies are available upon request. Other plasmids employed have been previously described in the corresponding published studies (see **Table S2**). The oligonucleotides used in this study are listed in **Table S3**.

### Polysome profile analysis

Cell extracts for polysome analysis were prepared according to previously established procedures [48]. Ten A_260_ units of cell extract were loaded onto 7-50% sucrose gradients and centrifuged in a Beckman Coulter SW41Ti rotor at 39,000 rpm for 2 h 45 min. Gradient fractionation profiles were obtained by continuously monitoring of the A_254_ using an ISCO UA-6 system. All steps of the procedure, except for profile acquisition, were performed at 4 °C.

### Fluorescence microscopy

To examine the localization of eL22 assembly, the JFY012 strain *(rpl22A*::kanMX4 *rpl22B*::kanMX4) was transformed with two plasmids: one expressing a yEGFP-tagged eL22A protein and another expressing the Nmd3-Δ100 protein variant, which dominantly inhibits the nucleo-cytoplasmic transport of LSUs [18, 49]. Cells were grown in liquid SD-Ura-Leu medium and subsequently shifted to SGal-Ura-Leu medium for 24 h. Following induction, cells were stained with Hoechst and visualized by fluorescence microscopy using an Olympus BX61 system. Acquired images were processed and analyzed with the Olympus cellSens software.

To investigate the subcellular localization of pre-ribosomal particles in the absence of eL22, the BY4741 and JFY012 strains were transformed with plasmids expressing Nop1-mRFP together with either uL23-yEGFP or uS3-yEGFP (see **Table S2**). Cells were grown in SD-Ura medium to mid-log phase, washed with PBS buffer (140 mM NaCl, 8 mM Na_2_HPO_4_, 1.5 mM KH_2_PO_4_, 2.75 mM KCl, pH 7.3) and subsequently examined by fluorescence microscopy.

### RNA extractions and analysis of steady-state RNA levels

Total RNA was extracted from samples corresponding to 10 OD_600_ units of mid-log phase cultures using the acid-phenol method [50]. Equal amounts of total RNA (typically 5 µg) were loaded and separated by electrophoresis on either 7% polyacrylamide-8 M gels for low molecular-weight RNAs or 1.2% agarose-6% formaldehyde gels for high molecular-weight RNAs. Following electrophoresis, RNAs were transferred onto nylon membranes and crosslinked as previously described [51]. For northern hybridization, specific antisense oligonucleotides (**see Table S3**) were 5’-end labelled with [γ-^32^P] ATP (6000 Ci/mmol; PerkinElmer) and used as probes to detect distinct pre-rRNAs and mature rRNAs. For primer extension analysis, equal amounts of total RNA were subjected to reverse transcription using the oligonucleotide probe g, end-labeled with [γ-^32^P] ATP, as primer. After primer extension, the resulting cDNAs were resolved by electrophoresis on 7% polyacrylamide-8 M sequencing gels. Gels were transferred onto Whatman filter papers and dried under vacuum. Nylon membranes or filter paper sheets were exposed to phosphorimager screens (Fuji BAS-IP MS; GE Healthcare) for variable times depending on signal intensity. Signals were visualized using a Typhoon™ FLA-9000 imaging system (GE Healthcare). When required, signal intensities were quantified using GelQuant.NET software (biochemlabsolutions.com).

### Pulse-chase labeling of pre-rRNAs

Pulse-chase labeling experiments were performed as previously described [52]. The BY4741 and JFY012 strains were first transformed with the empty plasmid YCplac33 to restore uracil prototrophy. Transformants were grown in liquid SD-Ura medium at 30 °C to mid-log phase, after which 40 OD_600_ units of cells were pulse-labelled for 2 min with 100 μCi of [5,6-^3^H]uracil (45-50 Ci/mmol, PerkinElmer). The pulse was terminated by the addition of an excess of unlabeled uracil (1 mg/ml), and cells were chased for 0, 5, 15, 30, and 60 min. Following the chase, cells were washed with cold water and flash-frozen in liquid nitrogen prior to RNA extraction. Total RNA was extracted using the hot acidic phenol-chloroform procedure described above. Incorporation of radiolabeled uracil was quantified using a scintillation counter, and approximately 3,000 cpm of each RNA sample were loaded and separated by electrophoresis on either a 7% polyacrylamide-8 M urea gels or a 1.2% agarose-6% formaldehyde gels. Following electrophoresis, RNA was transferred onto nylon membranes, crosslinked, and exposed to a ^3^H-sensitive screen (BAS-IP TR2040E; GE Healthcare) for variable exposure times. Signals were detected using a Typhoon™ FLA-9000 imaging system (GE Healthcare).

### Affinity purification of complexes containing eL22A-yEGFP

The JFY012 strain was transformed with either YCplac111-L22A-HA or YCplac111-L22A-GFP (**Table S2**). Untagged and yEGFP-tagged eL22 proteins were then immunoprecipitated using the established one-step GFP-Trap_A procedure (Chromotek) [53]. Approximately 40 OD_600_ units of mid-log phase cells grown in SD-Leu medium were harvested and washed twice with cold water. Cells were resuspended in 500 µl of lysis buffer (10 mM Tris-HCl, pH 7.5, 150 mM NaCl, 0.5 mM EDTA, 1.5 mM MgCl_2_, 1 mM DTT, 0.5% NP-40, and cOmplete™ Protease Inhibitor; Roche), and 300 µg of glass beads (0.45 µm diameter) were added. Cell lysis was performed using a FastPrep-24 homogenizer (MP Biomedicals), and lysates were clarified by centrifugation at maximum speed for 15 min.

Approximately 50 µl of GFP-Trap_A beads were added to each clarified extract and incubated for 2 h at 4 °C with end-over-end rotation. Beads were subsequently washed three times with 1 ml of lysis buffer and collected. RNA was purified using the phenol-chloroform extraction procedure and analyzed by northern hybridization as described above. RNA was also extracted from total cell lysates, and both immunoprecipitated and total RNA samples were included in the northern blot analyses.

## Data availability

All data presented in this study are included in this article, either in its main body or in its Supporting Information. Further information related to this study is available from the corresponding author upon request.

## Supporting information

Supplementary material to this article can be found online at https://XXX

## Acknowledgements

This article is dedicated to the memory of Prof. E. Cerdá-Olmedo. We thank all researchers cited in the manuscript for their generous gifts of materials.

## Funding

This publication is part of the R+D+I project PID2022-136564NB-I00, funded by MICIU/AEI/10.13039/501100011033 and ERDF/EU (“A way of making Europe”) to J.d.l.C. He also acknowledges support from the Andalusian Platform of Biomodels and Resources in Genome Editing, the FORTALECE Program (FORT23/00008) from MICIU, and the COST Actions Translacore (CA21154) and ProteoCure (CA20113) of the European Cooperation in Science and Technology. This research was also funded by National Institutes of Health R01 GM028301 to J. L. Woolford. J.F.-F. acknowledges an FPI fellowship from MCIN (BES-2017-080876).

## CRediT authorship contribution statement

**José Fernández-Fernández:** Methodology, Validation, Formal analysis, Investigation, Data curation, Writing-original draft, Writing-review & editing. **Sara Martín-Villanueva:** Methodology, Validation, Formal analysis, Investigation, Data curation, Writing-original draft, Writing-review & editing. **Taylor N. Ayers:** Methodology, Validation, Formal analysis, Investigation, Data curation, Writing-review & editing. **Carla V. Galmozzi:** Methodology, Formal analysis, Investigation, Writing-review & editing. **John L. Woolford:** Conceptualization, Methodology, Formal analysis, Resources, Writing-review & editing, Supervision, Funding acquisition. **Jesús de la Cruz:** Conceptualization, Methodology, Formal analysis, Resources, Writing-original draft, Writing-review & editing, Supervision, Project administration, Funding acquisition.

## Conflict of interest

All the authors declare that they do not have any conflicts of interest in relation to this work.

## Abbreviations

The abbreviations used are:

r-subunit: ribosomal subunit
rRNA: ribosomal RNA
r-protein: ribosomal protein
SSU: small or 40S ribosomal subunit
LSU: large or 60S ribosomal subunit
cryo-EM: cryo-electron microscopy
pre-rRNA: precursor rRNA

## Notes

### Competing Interest Statement

The authors have declared no competing interest.

