## Supplementary data for "Ribosomal protein eL22 contributes to the assembly of 60S ribosomal subunits in *Saccharomyces cerevisiae*"

### Supplementary Table S1. Yeast strains

| Strain | Relevant genotype | Source |
|---|---|---|
| BY4741 | MATa his3Δ1 leu2Δ0 met15Δ0 ura3Δ0 | Euroscarf |
| Y22672 | MATa/MATα his3Δ1/his3Δ1 leu2Δ0/leu2Δ0 met15Δ0/MET15 LYS2/lys2Δ0 ura3Δ0/ura3Δ0 RPL22A/rpl22A::kanMX4 | Euroscarf |
| Y25844 | MATa/MATα his3Δ1/his3Δ1 leu2Δ0/leu2Δ0 met15Δ0/MET15 LYS2/lys2Δ0 ura3Δ0/ura3Δ0 RPL22B/rpl22B::kanMX4 | Euroscarf |
| JFY010 | MATα his3Δ1 leu2Δ0 met15Δ0 ura3Δ0 rpl22A::kanMX4 | This work |
| JFY011 | MATa his3Δ1 leu2Δ0 met15Δ0 ura3Δ0 rpl22B::kanMX4 | This work |
| JFY012 | MATα his3Δ1 leu2Δ0 met15Δ0 ura3Δ0 rpl22A::kanMX4 rpl22B::kanMX4 | This work |
| Y25234 | MATa/MATα his3Δ1/his3Δ1 leu2Δ0/leu2Δ0 met15Δ0/MET15 LYS2/lys2Δ0 ura3Δ0/ura3Δ0 RPL38/rpl38::kanMX4 | Euroscarf |
| Y23772 | MATa/MATα his3Δ1/his3Δ1 leu2Δ0/leu2Δ0 met15Δ0/MET15 LYS2/lys2Δ0 ura3Δ0/ura3Δ0 RPL31A/rpl31A::kanMX4 | Euroscarf |
| JFF084 | MATa his3Δ1 leu2Δ0 ura3Δ0 rpl38::kanMX4 | This work |
| JFY449 | MATa his3Δ1 leu2Δ0 ura3Δ0 rpl31A::kanMX4 | This work |
| JFY531 | MATa his3Δ1 leu2Δ0 met15Δ0 ura3Δ0 rpl22A::kanMX4 rpl22B::kanMX4 rpl31A::kanMX4 | This work |
| JFY114 | MATa his3Δ1 leu2Δ0 met15Δ0 ura3Δ0 rpl22A::kanMX4 rpl22B::kanMX4 rpl38::kanMX4 | This work |
| Y23053 | MATa/MATα his3Δ1/his3Δ1 leu2Δ0/leu2Δ0 met15Δ0/MET15 LYS2/lys2Δ0 ura3Δ0/ura3Δ0 RPL19B/rpl19B::kanMX4 | This work |
| SMY714 | MATa his3Δ1 leu2Δ0 met15Δ0 ura3Δ rpl19B::kanMX4 | This work |
| SMY712 | MATα his3Δ1 leu2Δ0 met15Δ0 lys2Δ0 ura3Δ0 rpl22A::kanMX4 rpl22B::kanMX4 rpl19B::kanMX4 | This work |

### Supplementary Table S2. Plasmids

| Name | Relevant information | Source |
|---|---|---|
| YCplac33 | *CEN, URA3* | [1] |
| YCplac111 | *CEN, LEU2* | [1] |
| YCplac111-L22A-HA | *RPL22A-HA, CEN, LEU2* | This study |
| YCplac111-L22A-GFP | *RPL22A-yEGFP, CEN, LEU2* | This study |
| YCplac111-yEGFP | *CEN, yEGFP/TCYC1* | Kressler's lab |
| pRS316-RPL25-eGFP NOP1-mRFP | *RPL25-yEGFP, mRFP-NOP1, CEN, URA3* | [2] |
| pRS316-RPS3-eGFP NOP1-mRFP | *RPS3-yEGFP, mRFP-NOP1, CEN, URA3* | [2] |
| pRS316-GAL-*NMD3Δ100* | *GAL::NMD3Δ100, CEN, URA3* | [3] |

### Supplementary Table S3. Oligonucleotides

| Name | 5'-3' sequence | Use |
|---|---|---|
| L22A-UP | CTGCAGCCATTGACCTTATGTTCG | Cloning |
| L22A-DOWN | CTGCAGGAGTTATTAGGTCGAGGC | Cloning |
| L22A-GFP UP | GAATTCTGGGTTTGAAATCA | Cloning |
| L22A-GFP DOWN | TCTAGATTCTTCGTCTTCTTCTTCG | Cloning |
| L22A-COMP-UP | CAACCCAAAGACTTTGGAAT | PCR verification |
| L22A-COMP-DOWN | ACCATATAGCCATTTCGACT | PCR verification |
| L22B-COMP-UP | CCGTATCGTGCACAGATTTA | PCR verification |
| L22B-COMP-DOWN | AGGGCCAAATCTATGTACTG | PCR verification |
| L22A-SEC1 | GAGATTAACAACTTTACCTG | Sequencing verification |
| L22A-SEC2 | GATATAGACTTCCAGAAAGC | Sequencing verification |
| L22A-SEC3 | CTCTACCAAGACCAACGAAT | Sequencing verification |
| RPL31A-COMP-UP | GTCTGAGTAACTTGTGGAGTC | PCR verification |
| RPL31A-COMP-DOWN | GCGTACATCCTAGCCTAAC | PCR verification |
| RPL38-COMP-UP | GAATTCGCTCCCAGCAGATCCAAG | PCR verification |
| RPL38-COMP-DOWN | GGATCCGATGAATTTTTGAATGAGGCTCC | PCR verification |
| RPL19B -+1kb UP | CTCCAACCAGAGGTGCTGA | PCR verification |
| RPL19B -+1kb DW | TGGGCAATTCCAGTGAACGC | PCR verification |
| Probe b (18S) | CATGGCTTAATCTTTGAGAC | 18S rRNA hybridization |
| Probe c (3-D/$A_2$) | GACTCTCCATCTCTTGTCTTCTTG | Pre-rRNA hybridization |
| Probe d ($A_2$/$A_3$) | TGTTACCTCTGGGCCC | Pre-rRNA hybridization |
| Probe e (5.8S) | TTTCGCTGCGTTCTTCATC | 5.8S rRNA hybridization |
| Probe f (E/$C_2$) | GGCCAGCAATTTCAAGTTA | Pre-rRNA hybridization |
| Probe g ($C_1$/$C_2$) | GAACATTGTTCGCCTAGA | Pre-rRNA hybridization and primer extension |
| Probe h (25S) | CTCCGCTTATTGATATGC | 25S rRNA hybridization |
| Probe 5S | GGTCACCCACTACACTACTCGG | 5S rRNA hybridization |

### LEGENDS TO SUPPLEMENTARY FIGURES

**Figure S1. Pre-rRNA processing in *S. cerevisiae*.** (**A**) Schematic representation of an rDNA repeat unit. Each unit contains two independently transcribed elements. The long element is transcribed by RNA polymerase I (RNA pol I) into a polycistronic pre-rRNA encoding the 18S, 5.8S and 25S rRNAs. The short element is transcribed by RNA polymerase III (RNA pol III) into a pre-5S rRNA. External, internal, and non-transcribed spacers (ETSs, ITSs, and NTSs) are indicated. Mature rRNAs are shown as blue bars, and spacer regions as black lines. Transcription start sites are marked by red arrows. Processing sites and the locations of the probes used in this study (**Table S3**), are also indicated. (**B**) Schematic overview of the pre-rRNA processing pathway. The RNA pol I transcript undergoes co- or post-transcriptional processing, generating the 20S and 27SA$_2$ pre-rRNAs, which are incorporated into early pre-SSU and pre-LSU particles, respectively. These precursors are further processed into mature 18S, 5.8S, and 25S rRNAs by site-specific cleavages and 5'→3' or 3'→5' exonucleolytic trimming reactions. This figure has been adapted from that shown in García-Gómez *et al.* [4]. For additional details on the yeast pre-rRNA processing pathway, see references [5-7].

**Figure S2. Molecular environment of eL22 in the yeast LSU.** Position of eL22 and its surrounding molecular environment within the three-dimensional structure of the mature LSU from *S. cerevisiae*. Structural representations were generated using UCSF Chimera [8] based on the atomic model of the yeast 80S ribosome (PDB ID: 4V88; [9]). (**A**) View of the LSU from the inter-subunit interface and after a 90 º rotation around the X-axis. R-proteins from 25S rRNA domain III are highlighted by surface rendering. Other r-proteins are shown in light blue, and the 25S, 5S, and 5.8S rRNAs are displayed in pale grey. (**B**) Close-up view of r-proteins bound to 25S rRNA domain III. The surface of each r-protein is shown in a distinct color, whereas the rRNA is colored purple. Views correspond to those shown in panel A.

**Figure S3. Functional characterization of the eL22A-yEGFP construct.** (**A**) Growth comparison of isogenic wild-type and *rpl22*Δ strains transformed with an empty centromeric YCplac111 plasmid, and of the *rpl22*Δ strain transformed with a YCplac111 plasmid harboring an RPL22A-yEGFP allele that expresses a C-terminal yEGFP-tagged eL22A from its endogenous promoter. Cells were grown in liquid SD-Leu medium at 30 °C to mid-log phase, diluted to an $OD_{600}$ of 0.05, and 10-fold serial dilutions were spotted onto SD-Leu plates. Plates were incubated for 2 days at 30 °C. (**B**) Polysome profile analysis of the strains shown in panel A. The above strains were grown in liquid SD-Leu medium until mid-log phase at 30 °C. Whole-cell extracts were prepared from mid-log-phase cultures, and 10 $A_{260}$ units of each extract were resolved in 7-50% sucrose gradients. The $A_{254}$ was continuously recorded. Sedimentation is from left to right. Peaks corresponding to free SSUs (40S), free LSUs (60S), vacant ribosomes/monosomes (80S), and polysomes are indicated. Half-mer polysomes are marked with arrows.

**Figure S4. Nucle(ol)ar assembly of eL22.** Subcellular localization of GFP-tagged eL22 upon induction of the dominant-negative Nmd3-Δ100 protein, expressed from a *GAL* promoter. Cells of the *rpl22* null strain expressing eL22-yEGFP were transformed with pRS316-GAL-NMD3Δ100, grown in SD-Ura-Leu medium (Glc) at 30 °C to mid-log phase, and shifted to SGal-Ura-Leu (Gal) for 24 h to induce Nmd3-Δ100 expression. Cells were stained with Hoechst, and yEGFP and Hoechst signals were examined by fluorescence microscopy. Arrows indicate nuclei of representative cells. Approximately 200 cells were analyzed per experiment; the percentage of cells showing nucle(ol)ar accumulation of eL22-yEGFP is indicated (ca. 80%). Scale bar, 10 µm.

**Figure S5. The LSU deficit caused by loss of eL22 is attenuated at high temperatures.** Polysome profile analysis of isogenic wild-type and *rpl22*Δ strains grown in liquid YPAD medium to mid-log phase at 22 °C (**A**) or 37 °C (**B**). Whole-cell extracts were prepared, and 10 $A_{260}$ units of each extract were resolved in 7-50% sucrose gradients. The $A_{254}$ was continuously recorded. Sedimentation is from left to right. Peaks corresponding to free SSUs (40S), free LSUs (60S), vacant ribosomes/monosomes (80S) and polysomes are indicated. Half-mer polysomes are marked with arrows.

**Figure S6. Pre-rRNA processing is largely unaffected in *rpl22* null cells grown at elevated temperatures.** Isogenic wild-type and *rpl22*Δ strains were grown in liquid YPAD medium at 22, 30 and 37 °C to mid-log phase. Total RNA was extracted, and equal amounts (5 µg) were analyzed by northern blotting. The same membrane was sequentially hybridized with the probes indicated in parentheses and described in Table S3. Signal intensities of the different RNAs were quantified by phosphorimager scanning (values shown below each panel) and normalized to wild-type levels, arbitrarily set to 1.0.

**Figure S7. eL22 is required for efficient nuclear-cytoplasmic export of pre-LSU particles.** Wild-type and *rpl22*Δ strains were transformed with plasmids co-expressing mRFP-Nop1 and either uL23-yEGFP or uS3-yEGFP from their endogenous promoters. The uL23 r-protein was used as a reporter for pre- and mature LSU particles, whereas the uS3 r-protein served as a reporter for pre- and mature SSU particles. Cells were grown in liquid SD-Leu medium at 30 °C to mid-log phase and shifted to 22 °C for 3 h. Subcellular localization of the yEGFP-tagged r-proteins and the mRFP-Nop1 nucleolar marker was analyzed by fluorescence microscopy. Selected nucleoli are indicated by arrows. Approximately 200 cells were examined per experiment; the percentage of cells showing nucle(ol)ar accumulation of each reporter is indicated. Scale bar, 10 µm.

**Figure S8. Ribosome-targeting drugs exacerbate the growth defect of cells lacking eL22 and eL38 r-proteins.** Wild-type, *rpl38*Δ, *rpl22*Δ and *rpl38*Δ *rpl22*Δ strains were grown in liquid YPAD media at 30 °C to mid-log phase, diluted to an $OD_{600}$ of 0.05, and 10-fold serial dilutions were spotted onto YPAD plates lacking or containing sublethal concentrations of cycloheximide (0.075 µg/ml) or neomycin (5 mg/ml). Plates were incubated at 30 °C for 3 days (YPAD and cycloheximide-containing plates) or 4 days (neomycin-containing plates).

### SUPPLEMENTARY FIGURES

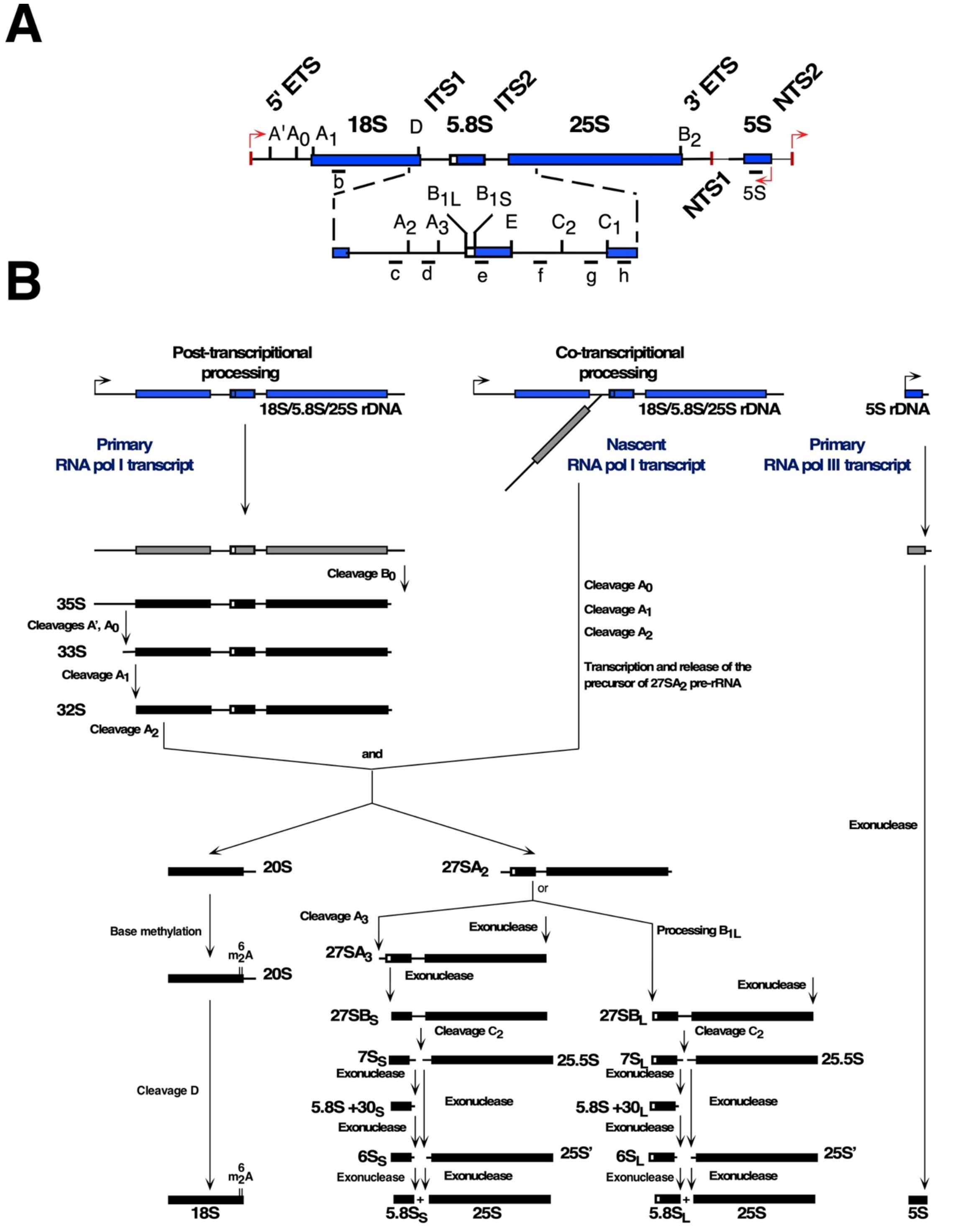


**Figure S1**

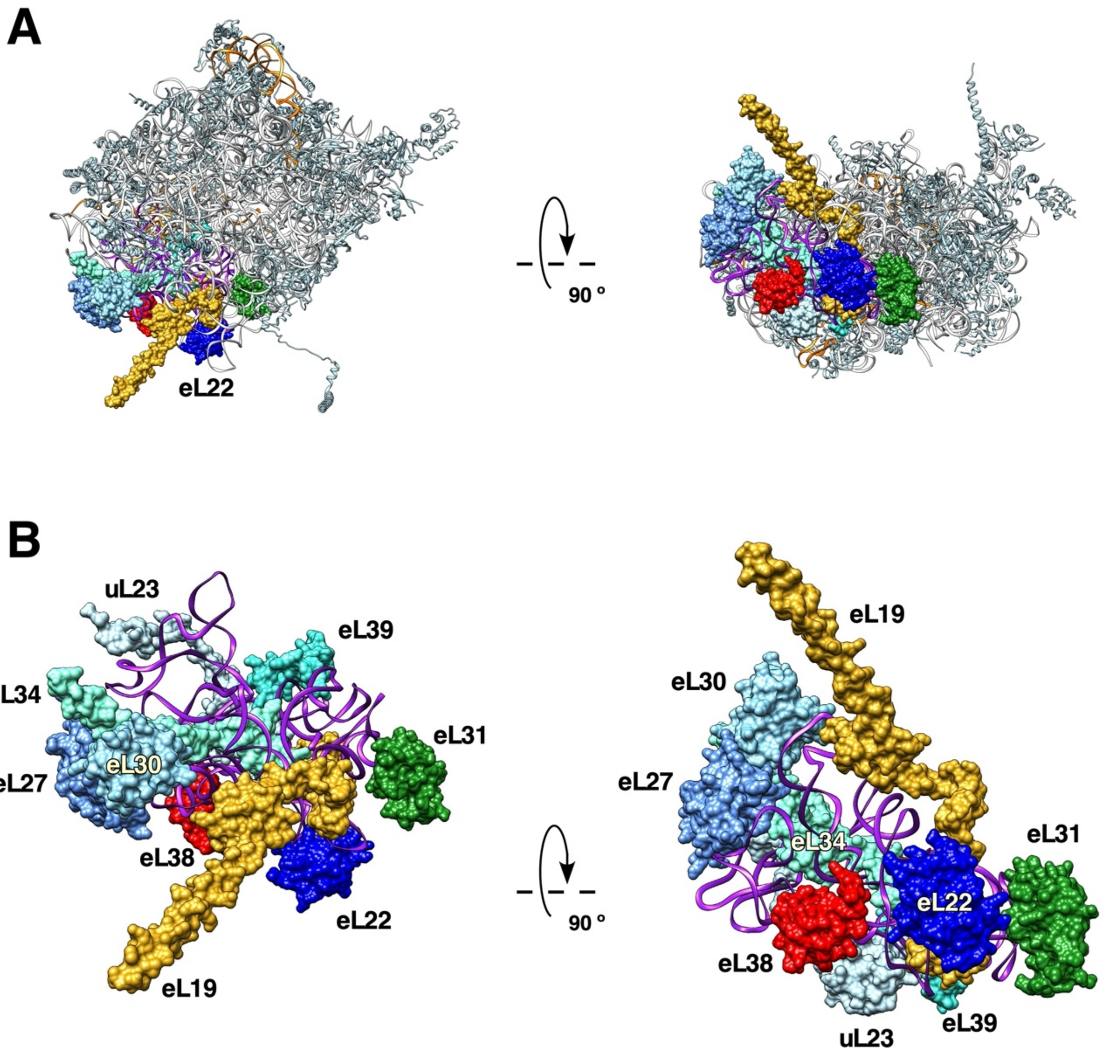
A
eL22
90 °
B
uL23
eL39
eL34
eL31
eL30
eL27
eL38
eL22
eL19
eL19
eL30
eL27
eL31
eL34
90 °
eL22
eL38
eL39
uL23

Figure S2

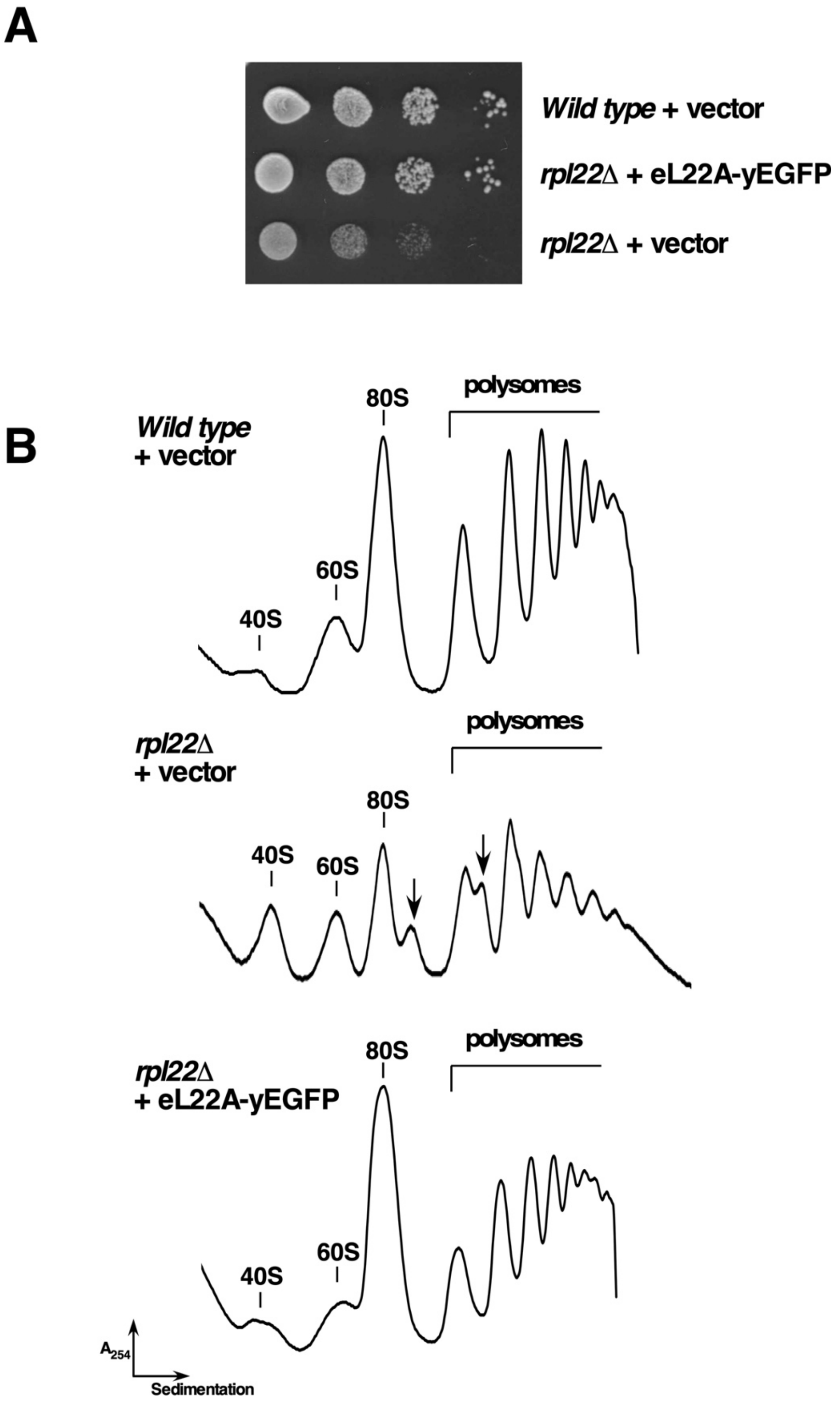
A
Wild type + vector
rpl22Δ + eL22A-yEGFP
rpl22Δ + vector
B
Wild type
+ vector
polysomes
80S
60S
40S
rpl22Δ
+ vector
polysomes
80S
40S
60S
rpl22Δ
+ eL22A-yEGFP
80S
polysomes
60S
40S
A254
Sedimentation

**Figure S3**

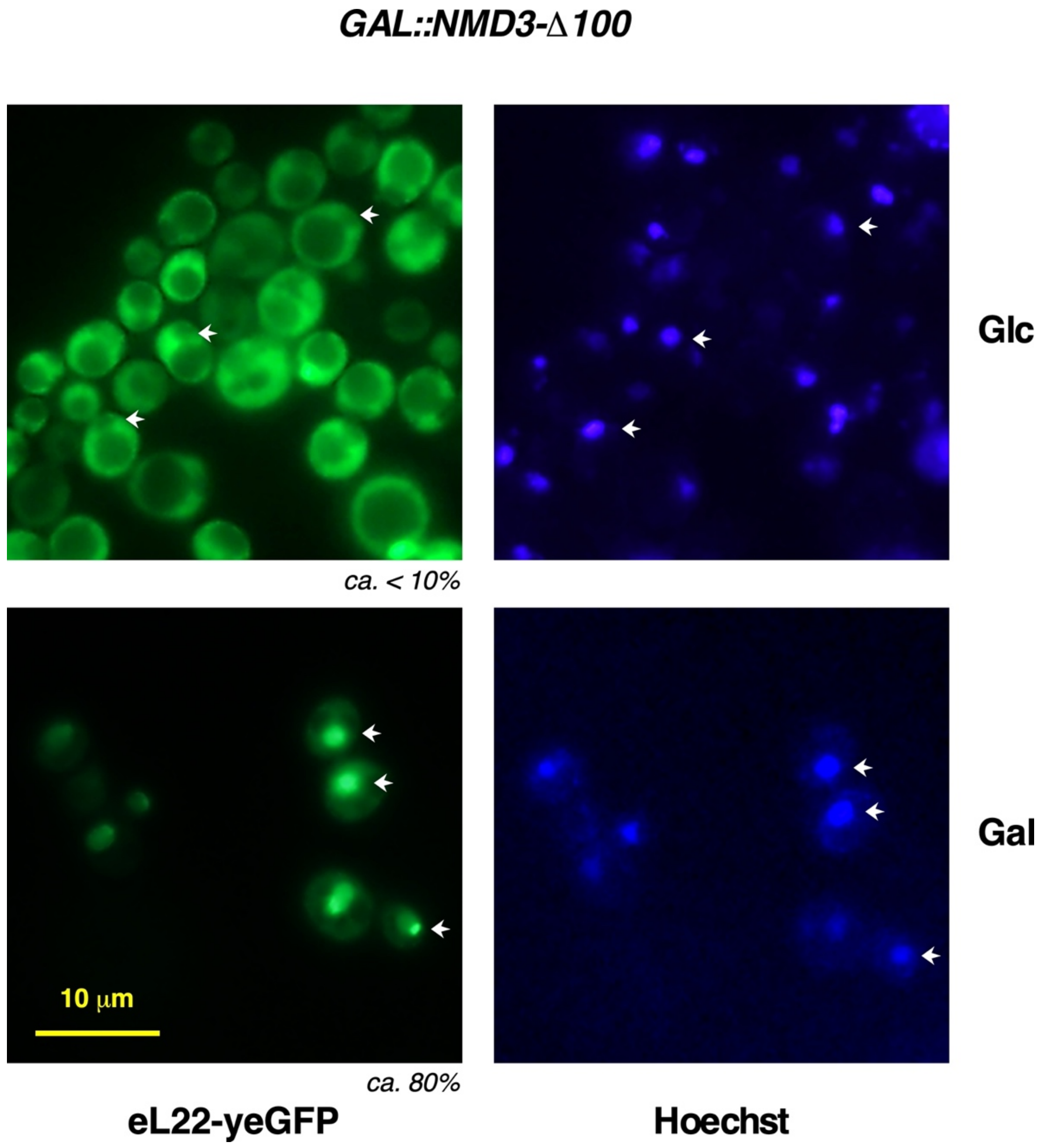


**Figure S4**

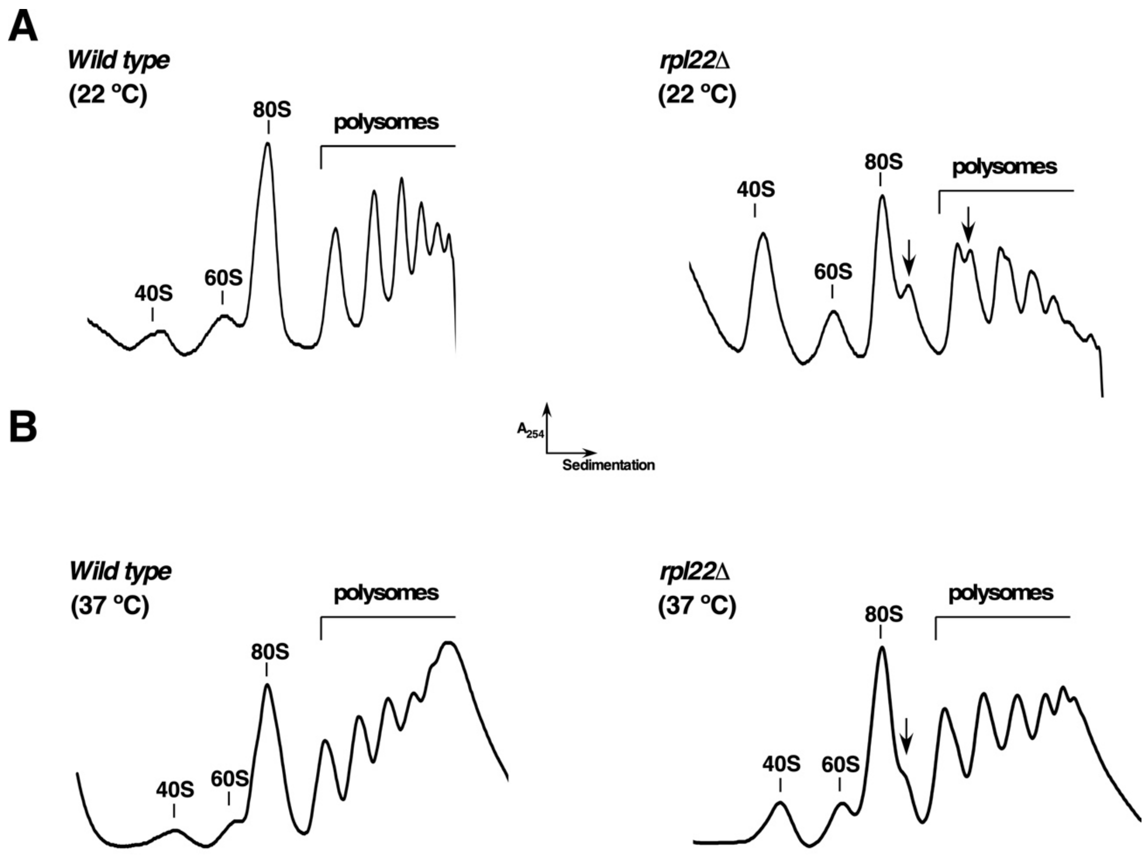
A
Wild type
(22 °C)
80S
polysomes
40S
60S
rpl22Δ
(22 °C)
40S
80S
polysomes
60S
B
A254
Sedimentation
Wild type
(37 °C)
polysomes
80S
40S
60S
rpl22Δ
(37 °C)
80S
polysomes
40S
60S

**Figure S5**

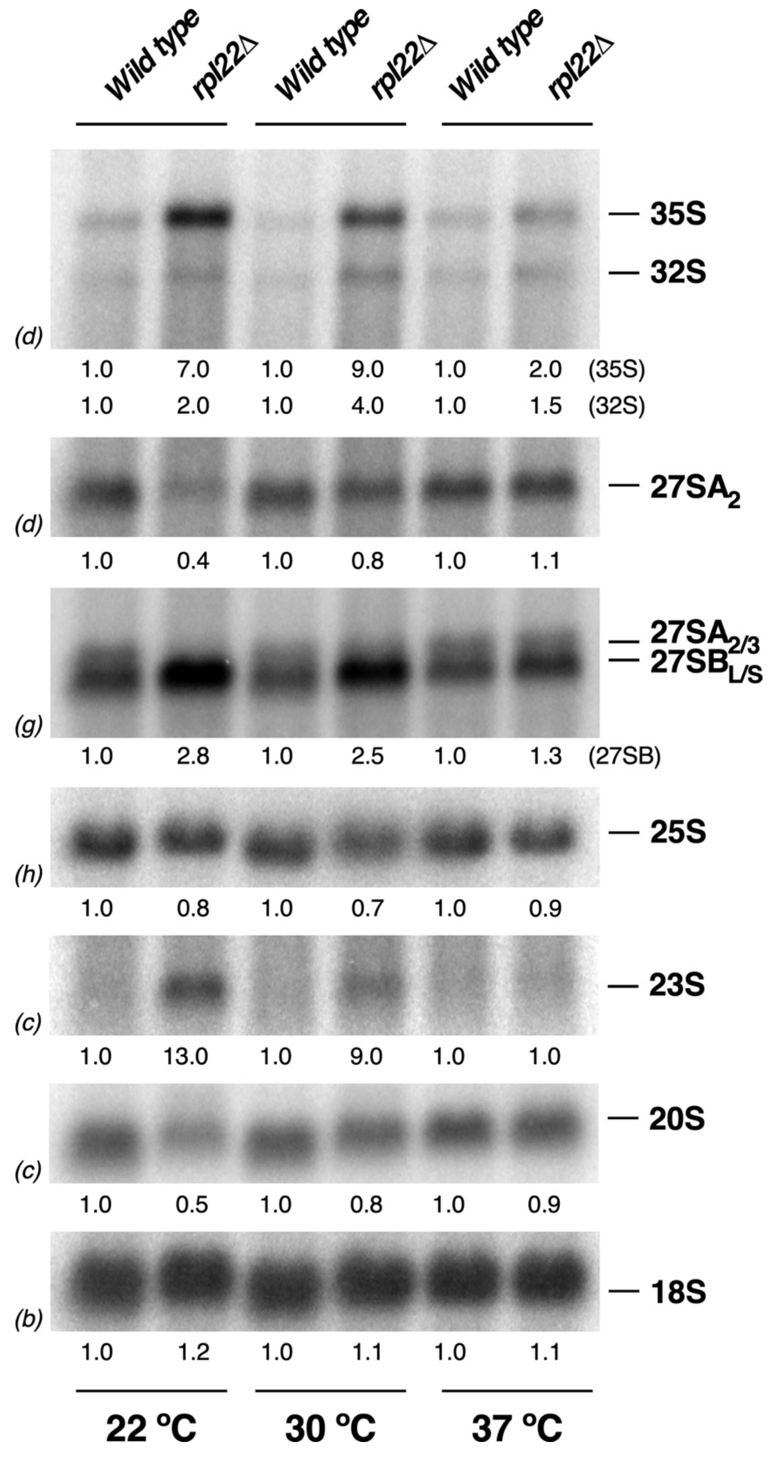
Wild type
rpl22Δ
Wild type
rpl22Δ
Wild type
rpl22Δ
— 35S
— 32S
(d)
1.0 7.0 1.0 9.0 1.0 2.0 (35S)
1.0 2.0 1.0 4.0 1.0 1.5 (32S)
— 27SA2
(d)
1.0 0.4 1.0 0.8 1.0 1.1
— 27SA2/3
— 27SBL/S
(g)
1.0 2.8 1.0 2.5 1.0 1.3 (27SB)
— 25S
(h)
1.0 0.8 1.0 0.7 1.0 0.9
— 23S
(c)
1.0 13.0 1.0 9.0 1.0 1.0
— 20S
(c)
1.0 0.5 1.0 0.8 1.0 0.9
— 18S
(b)
1.0 1.2 1.0 1.1 1.0 1.1
22 ºC
30 ºC
37 ºC

**Figure S6**

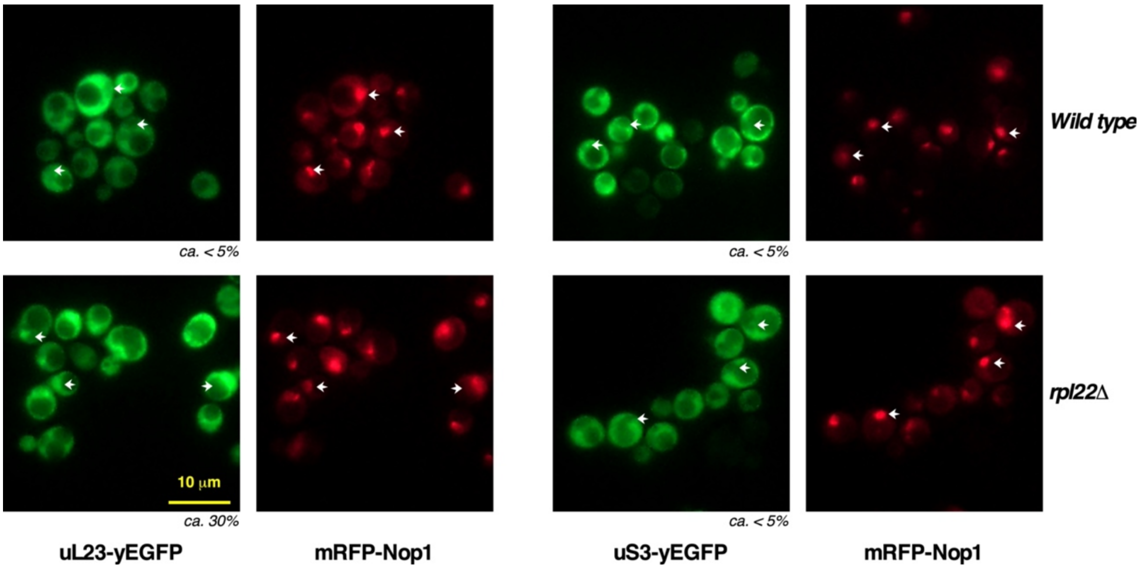
Wild type
rpl22Δ
ca. < 5%
ca. < 5%
10 μm
ca. 30%
ca. < 5%
uL23-yEGFP
mRFP-Nop1
uS3-yEGFP
mRFP-Nop1

**Figure S7**

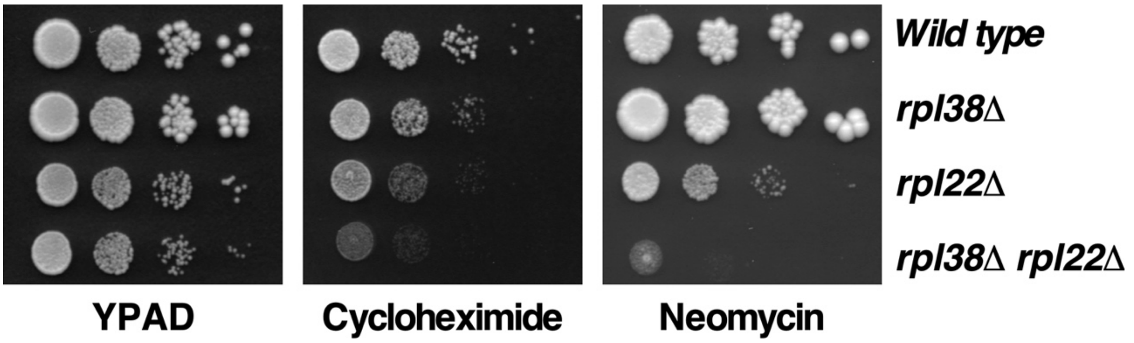
Wild type
rpl38Δ
rpl22Δ
rpl38Δ rpl22Δ
YPAD
Cycloheximide
Neomycin

**Figure S8**
